# Ultra-small bispecific fusion protein augments tumor penetration and treatment efficacy for pancreatic cancer

**DOI:** 10.1101/2022.05.17.492287

**Authors:** Qian Wang, Jingyun Wang, Hao Yan, Zheng Li, Kun Wang, Feiyu Kang, Jie Tian, Xinming Zhao

**Author notes:** These authors contributed equally to this work (Qian Wang, Jingyun Wang).

## Abstract

Pancreatic ductal adenocarcinoma (PDAC) is one of the leading causes of cancer-related deaths. Due to drugs’ low intrinsic anticancer activity and the unique physiological barrier of PDAC tumors, the once highly anticipated antibody-based pathway-targeted therapies have not achieved promising improvement in outcomes. Here, an ultra-small-sized bispecific fusion protein, termed Bi-fp50, that could largely enrich deep tumor tissue and effectively inhibit PDAC tumor growth was reported. The bispecific Bi-fp50 protein was constructed by a typical synthetic biology method that could efficiently target EGFR and VEGF of PDAC cells simultaneously *in vitro* and *in vivo*. For Bxpc3 and Aspc1 PDAC cells, the Bi-fp50 achieved a significant and synergistic therapeutic effect. Owing to the small size of only 50 kDa and the function of reducing the interstitial fluid pressure by vascular normalization, the Bi-fp50 showed enhanced penetration, considerable accumulation, and uniform distribution in tumor and subsequently led to effective inhibition of the growth of Bxpc3 cells-induced PDAC tumor *in vivo*. Furthermore, no noticeable side effect of Bi-fp50 was found *in vitro* and *in vivo*. This work demonstrates that the synthetic Bi-fp50 fusion protein could be used as a new effective pathway-specific targeted therapy for PDACs.

## Introduction

Pancreatic cancer therapy currently remains a formidable challenge. The overall 5-year survival for pancreatic cancer, mainly comprising pancreatic ductal adenocarcinoma (PDAC), has changed little over the past decade [1-2]. Surgery, adjuvant or neoadjuvant chemotherapy and radiotherapy, nab-paclitaxel–gemcitabine, and FOLFIRINOX-based chemotherapy strategies are the standard treatment options in patients in different stages of PDAC [3-5]. Although these treatments can significantly prolong the survival of patients, they also lack high targeting and specificity, which would dramatically affect the therapeutic effect, while the PDACs have an exceptionally high molecular heterogeneity [6-7]. Numerous mono-targeted agents have been evaluated alone or combined with chemotherapy in PDAC treatment [8-9]. Unfortunately, most of these agents have failed to improve the patient survival from the clinic feedback [10-11]. It was speculated that the futility was significant because of the low intrinsic efficacy of drugs and the unique physiological barrier of PDAC tumors [3, 12-13]. Multi-specific therapeutic agents that work by engaging two or more entities are believed to lead the current fourth wave of biopharmaceutical innovation [14-15] and have shown enhanced or even synergistic therapeutic effects recently [16-17]. Therefore, the strategy of developing highly multi-specific therapeutic agents is promising to improve the targeted therapy of PDAC.

Epidermal growth factor receptor (EGFR) is overexpressed in more than 90% of patients’ PDAC, and signaling via it could regulate the growth, proliferation, and apoptosis of PDACs [13, 18]. Vascular endothelial growth factor receptor (VEGF) is another overexpressed target in 64% of PDAC and has been recognized as a critical mediator of angiogenesis to support tumorigenesis [19]. Interestingly, VEGF and EGFR share common downstream signaling pathways, and EGFR activation can drive VEGF gene expression, subsequently providing a positive feedback loop [20-21]. Substantial preclinical studies and multiple clinical trials in several cancer diseases all suggest that the EGFR and VEGF pathways are interrelated, and dual EGFR-VEGF pathway inhibition could delay acquired resistance of anti-EGFR or anti-VEGF used alone [22-24]. Therefore, it’s reasonable to hypothesize that dual inhibition of EGFR-VEGF pathways of PDAC cells simultaneously may delay the individual resistance and achieve an enhanced and synergistic therapeutic effect, which has not been achieved for PDACs according to our knowledge.

Furthermore, the unique physiological barriers of PDAC tumors, especially the extremely dense desmoplastic stroma and the increased interstitial fluid pressure, are one of the decisive factors affecting the therapeutic efficacy of the drug [3, 12-13, 25]. One effective strategy is treating tumor vasculature by targeting VEGF, which can gradually develop a “vessel-normalization window” and reduce the tumoral interstitial fluid pressure (IFP) [26-27]. Another strategy is to construct an ultrasmall-sized therapeutic agent, which was founded critical for overcoming pathophysiological barriers to efficient tumor penetration and accumulation [13, 28-30].

Herein, we developed an ultra-small bispecific fusion protein, termed Bi-fp50, that could target EGFR-VEGF pathways simultaneously for PDACs and efficiently extravasate through the dense barriers to accumulate in deep tumor tissue, subsequently achieving enhanced therapeutic effect for PDAC tumors (Scheme 1). The obtained bispecific therapeutic agent Bi-fp50 with a molecular weight of only 50 kDa was constructed by a typical synthetic biology method that reduced the molecular weight several times two corresponding antibodies. The synthesized Bi-fp50 agent showed excellent bispecific targeting for VEGF and EGFR and synergistic anticancer activity for PDACs *in vitro*. The anti-VEGF function of Bi-fp50 could normalize tumor vasculature, along with its ultra-small size, allowing the Bi-fp50 to penetrate dense barriers and enrich the whole Bxpc3 pancreatic tumor. After intravenous injection, the Bi-fp50 showed a significant tumor inhabitation effect in Bxpc3 cells-induced tumor *in vivo*, while no apparent side effects were found *in vitro* and *in vivo*. Our approach of constructing ultra-small-sized multi-specific protein provides a new strategy for more effective targeted pathway therapy for PDAC.

**Scheme 1.**
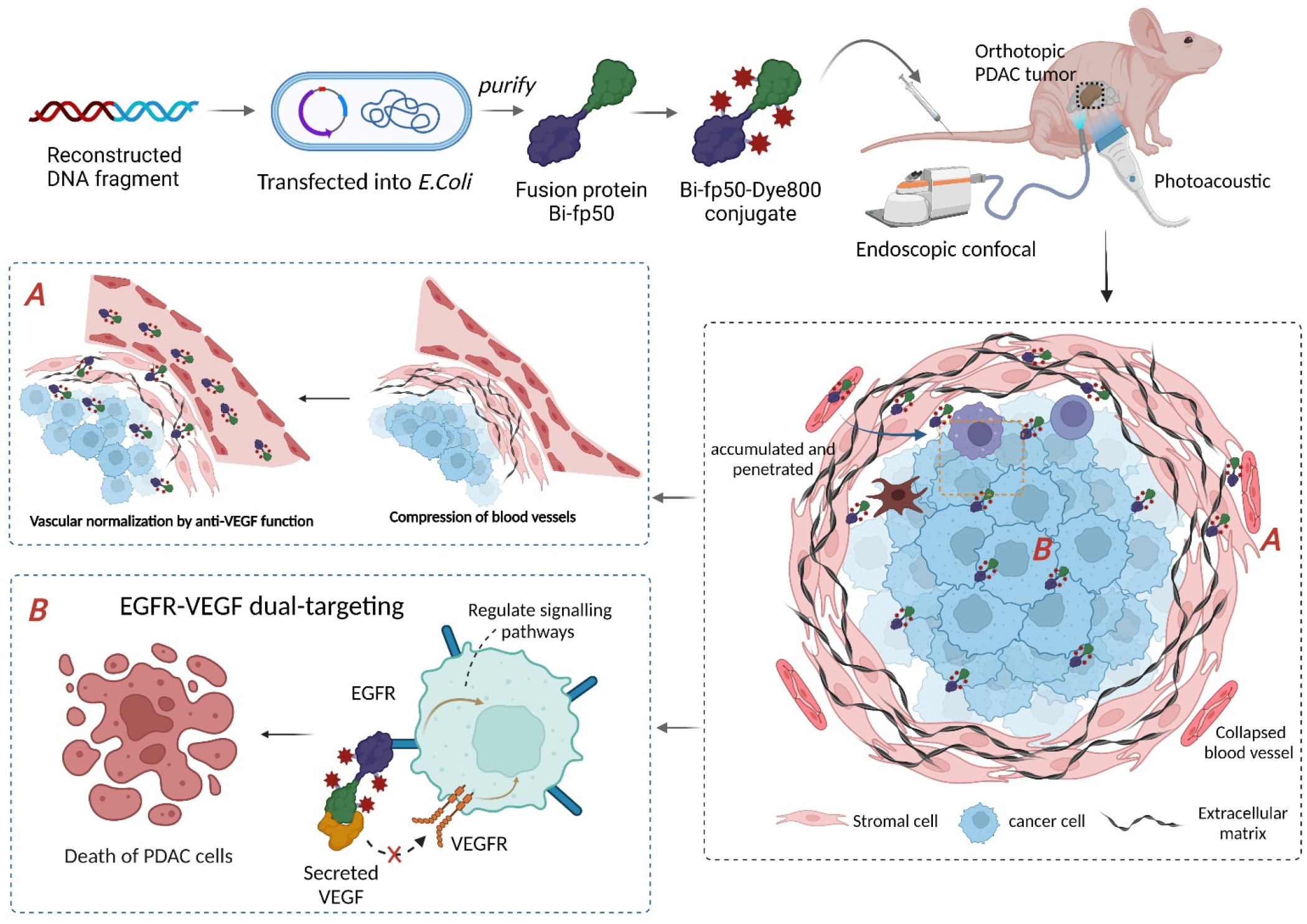
Schematic illustration of the construction of the bispecific fusion protein Bi-fp50 and its application for enhanced tumor penetration and targeted therapy against pancreatic cancer. The anti-VEGF function of Bi-fp50 could normalize the vessel structure of the tumor and reduce interstitial fluid pressure, along with its ultra-small size together enhance intra-tumoral penetration and accumulation. In addition, dual targeting of EGFR and VEGF simultaneously produced apoptosis of PDACs.

## Materials and methods

### Preparation of bispecific fusion protein Bi-fp50

The anti-VEGF scFv (selected) and anti-EGFR scFv (selected) with good affinity and specificity were screened and selected from the homemade human single-chain antibody library through commercial VEGF and EGFR proteins (R&D Systems, Inc.). Then, according to the connecting sequence of VH_1_-G_4_S-VL_1_-G_4_S-VL_2_-G_4_S-VH_2_ of the two selected scFvs, the corresponding gene sequence was constructed and expressed with the pET30a expression vector (Beijing Qingke Biotechnology Co. Ltd.). The genes of two VH CDR3 fragments were placed at both ends of the sequence to maintain the binding ability. Standard DNA sequencing analysis was used to verify that the gene sequence was correct and cloned in the frame. This obtained gene was then transformed into Escherichia coli strain BL21 (λDE3) to obtain the bispecific fusion protein Bi-fp50. After further purification by Ni-nitrilotriacetic acid (NTA), Bi-fp50 was used for further experiments. In addition, scFv2 directly connected anti-EGFR scFv with anti-VEGF scFv, and Bi-fp50x, a non-functional protein with a similar molecular weight to Bi-fp50, were also produced as the control groups.

### Characteristics of Bi-fp50

Sodium dodecyl sulfate-polyacrylamide gel electrophoresis (SDS-PAGE) was used to study the molecule weight of Bi-fp50, anti-EGFR scFv, and anti-VEGF scFv. And the Fourier transform infrared spectra (FTIR) with an FTIR NETZSCH X70 was used to characterize the structure of Bi-fp50. Meanwhile, the enzyme-linked immunosorbent assay (ELISA) assay was carried out to compare the binding affinities of Bi-fp50 compared with anti-EGFR scFv, anti-VEGF scFv, scFv2, and Bi-fp50x. First, EGFR or/and VEGF were coated on 96-well plates at 4 °C for 24 hours. The plates were then incubated with either anti-EGFR scFv, anti-VEGF scFv, scFv2, Bi-fp50, or Bi-fp50x at 37 °C for 2 hours. After reaction with 3,3’,5,5’-tetramethylbenzidine (TMB), the binding affinity of these proteins were identified by testing the absorbance at 450 nm. Rabbit mono/polyclonal antibodies against EGFR, VEGF, GAPDH, STAT3, p-STAT3, AKT, p-AKT, as well as β-actin were purchased from the Abcam platform for western blotting analysis of Bi-fp5o following typical standard protocol.

### Cell culture and *in vitro* targeting experiments

The human pancreatic ductal adenocarcinoma cell lines Bxpc3, Aspc1, Panc1, SW1990, Capan2, and T3M4, and HPDE6-C7 human pancreatic duct epithelial cells were purchased from ATCC (American Type Culture Collection). All cell lines were cultured in Dulbecco’s modified Eagle medium (DMEM) (HyClone) with 10% fetal bovine serum FBS (HyClone) and 1% penicillin-streptomycin (P.S.) (Promega) at 37 °C in a 5% CO_2_ atmosphere.

EGFR and VEGF expression for different PDAC cells were determined by Western blotting assay. Six kinds of pancreatic cancer cells were harvested during the exponential phase and lysed in 1×PBS, 1% Nonidet P-40 (Thermo Scientific), 2 μg ml^-1^ Aprotinin, 2 μg ml^-1^ Leupeptin, and 50 μg ml^-1^ phenylmethylsulfonyl fluoride (PMSF). After centrifuging at 12,000 g at 4 °C for 10 min, the total proteins were extracted from the supernatants and then quantified using the Pierce BCA assay kit (Thermo Scientific). Proteins from each cell line were separated through SDS-PAGE (10% resolving gel) and electroblotted onto a polyvinylidene fluoride (PVDF) membrane using the semi-dry blotting system (Bio-rad). Next, the membrane was blocked with 1× PBST buffer for 30 min, incubated with anti-EGFR and anti-VEGF (Abcam) at a concentration ratio of 1:1,000, respectively. And then anti-rabbit secondary antibodies at 1:2,000 conjugated with horseradish peroxidase. Chemiluminescence signals were detected using ImageQuant LAS 4000 (G.E. Healthcare Life Sciences).

The targeting properties of Bi-fp50 were further studied. The Bi-fp50, scFvs, and two anti-VEGF scFv and Bi-fp50x were labeled with Alexa Fluor 488 (AF488) separately to visualize the protein. After incubating different kinds of protein probes (0.05 μM) for 2 h, Bxpc3 cells were fixed and stained by DAPI (4’, 6-diamidino-2-phenylindole). Confocal microscopy was used to image the cellular targeting of protein probes. Image J software was further used for quantitative analysis.

### *In vitro* anticancer ability assessment of Bi-fp50

Two different cell lines (Bxpc3 and Axpc1) were used to study the anticancer effect of Bi-fp50. HPDE6-C7, Bxpc3, and Axpc1 cells were seeded on 96-well plates and cultured for another 24 hours. Then, anti-EGFR scFv, anti-VEGF scFv and Bi-fp50 with different concentration (0, 0.2, 0.3, 0.45, 0.6, 1.0 μM) were added into the culture plates and co-incubated at 37°C for another 24 h respectively. For cell viability assay, MTT (3-(4,5-Dimethylthiazol-2-yl)-2,5-Diphenyltetrazolium Bromide) solution was added to each well and incubated for 4 h at 37°C. Then, the cell viability was determined by detecting the absorbance at 570 nm.

For live/dead staining assay, 24 hours after being co-incubated with a different protein, the Bxpc3 cells were stained by Calcein–AM and propidium iodide (PI); a fluorescent inverted microscope system imaged qualitative staining results. The experiment was performed in triple technical replicates and standardized.

For apoptosis analysis, the Bi-fp50 and other control proteins were first co-incubated with Bxpc3 cells for 24 hours. Then the cells were stained by an annexin V-FITC apoptosis detection kit (Beyotime). Briefly, the cells were suspended in 5 µL of annexin V-FITC and 195 µL of annexin V-FITC binding buffer and incubated for 10 min. After centrifuging for 5 min, the cells were re-suspended in 190 µL of binding buffer and 10 µL of propidium iodide (P.I.) working solution. The flow cytometric method was finally carried out to analyze the staining results.

### Animals and PDAC tumor model

Animal experiments procedures were performed in compliance with the Animal Ethics Committee of the Chinese Academy of Medical Sciences Tumor Hospital (#NCC2019A010). The whole study maintained strict adherence to protocols for animal care and use. Bxpc3-Luc (5 × 10^5^ cells/mouse) cells were injected subcutaneously for the female BALB/C nude mice six weeks old (Weitong Lihua Experimental Animal Technology Co. Ltd. China). When subcutaneous tumors reached 5 mm, they were harvested and then chopped into one mm^3^ piece. And the small tumor masses were *in-situ* implanted on the pancreas to form the orthotopic PDAC tumor model (for imaging) or injected into the subcutaneous to form the subcutaneous PDAC tumor model (for therapy).

### Blood clearance, biodistribution, and intra-tumoral penetration of Bi-fp50 *in vivo*

Bi-fp50 (or scFv2) was labeled with IRDye 800CW dye (Dye 800, LI-COR Biotechnology) in the biodistribution and pharmacokinetics analysis. In the blood clearance measurement, 10 μl of blood was harvested from the tail vein of Bxpc3-Luc tumor-bearing mice post-injection of Bi-fp50-Dye800 at different time points. The blood sample was immediately dissolved in 1 ml lysis buffer. And the concentration of the Bi-fp50-Dye800 in the blood was measured by the fluorescence spectrum with the Ex/Em at about 800 nm/820 nm. A series of probe-blood dilutions were performed to obtain the standard calibration curve. Finally, a blank blood sample without Bi-fp50-Dye800 was measured to determine the blood autofluorescence level.

*In vivo* fluorescence imaging, Bxpc3-Luc tumor-bearing mice were intravenously injected with Bi-fp50-Dye800 or scFvs-Dye800 separately (2 mg Kg-1), and fluorescent signals were fluorescent detected at various time points by an IVIS imaging system (PerkinElmer). For the quantitative biodistribution analysis of protein in major metabolism organs and tumors, Bxpc3 tumor-bearing mice were sacrificed at 4, 8, and 12 hrs. Post-injection of probes. The major organs and tumors were weighed and homogenized in the lysis buffer and diluted 100 times for further fluorescence measurement. By measuring the fluorescence intensity of the probe in the tissue solution and comparing it with the standard curve, the concentration and amount of the probe in the tissue can be known. The samples were measured four times to ensure reproducibility and accuracy.

Intra-tumoral penetration of Bi-fp50 and scFv2 were further studied by a multispectral photoacoustic tomography system (iTheraMedical). The penetration and distribution of Bi-fp50 and scFv2 in the orthotopic PDAC tumor were analyzed using the 3D model of the photoacoustic imaging (PAI) device, and the results were recorded under different hours’ intervals post i.v. Injection. The ImageJ software was used to analyze the fluorescence intensity of Bi-fp50 quantitatively.

### Real-time monitoring of the dynamics of Bi-fp50 in PDAC tumor

The further intratumor penetration and dynamic distribution of Bi-fp50 in local tumor tissue were monitored by a fibered confocal fluorescence microscopy (FCFM) imaging system (Cellvizio). The orthotopic Bxpc3 tumor-bearing mice were intravenously injected with Bi-fp50-AF647 or scFv2-AF647 separately. FITC dyes were used to stain out the outline of the tissue. After anesthetized by peritoneal perfusion, a small incision was made in the left abdominal rectus of the mouse. A flexible fiber mini probe connected to a 488/660 nm laser scanning unit was then planted through the incision. To obtain high-resolution images, the mini probe was attached to the surface of normal and tumor tissues. Dynamic observations of the distribution of Bi-fp50-AF647 and scFv2-AF647 were visualized at 660 nm mode. The appropriate signal could be obtained by the gentle movement of the probe. The ImageJ software was used for quantitative analysis of the fluorescence intensity. After imaging, the tumors were harvested and fixed in 4% paraformaldehyde for 2 h at 4 °C and dehydrated in 15% sucrose for 4 h, followed by 25% sucrose for another 24 h. Then, the tumor tissues were embedded and cut into ∼10 μm cryosections. The slice was further rehydrated with PBS, blocked with 5% bovine serum albumin, and stained with DAPI to stain the nucleus.

### Antitumor effect of Bi-fp50 *in vivo*

10 mice with subcutaneous Bxpc3 tumors were randomly divided into two groups (Bi-fp50 and scFv2). After intravenously injected with Bi-fp50 (10 mg Kg^-1^) or scFv2 (10 mg Kg^-1^), tumor size and the bodyweight of mice were continually recorded every 2 days. The mice were sacrificed 21 days after treatment, and major organs were harvested, fixed in 4% neutral buffered formalin, embedded in paraffin, and sectioned for further histological analysis. To evaluate in-depth intra-tumoral treatment effects, another 6 mice with Bxpc3 tumors were euthanized 3 days after different treatments (Bi-fp50 and scFv2, n=3). The whole tumors were collected and fixed in 4% PFA for 1 day. Then the Bxpc3 tumor tissues were transferred for TUNEL staining following the standard protocol.

### Statistical Analysis

All quantitative data were recorded as mean ± standard S.D. Means were compared using Paired t-test and two-way analysis of variance (ANOVA). The Pearson test used the linear and non-linear tests. *p<0.05 were considered as statistical significance; **p<0.01, remarkably; ***p<0.001, very prominently.

## Results and discussion

### Synthesis and characteristics of Bi-fp50

The bispecific fusion protein Bi-fp50 was designed and synthesized by a typical synthetic biology strategy (Scheme 1). As with the previous procedure [30], we first screened and selected the anti-VEGF and anti-EGFR antibodies with high specificity to PDAC cells from a natural antibody library. Then, the corresponding gene sequence, which can express single-chain variable fragments (scFvs) of anti-EGFR and anti-VEGF, was constructed and transfected into engineered bacteria *E. Coli* by genetic engineering method (Fig. S1). After lysis and purification, the Bi-fp50 was obtained. Meanwhile, the control group protein scFv2 was synthesized by directly connecting the two scFv (anti-EGFR and anti-VEGF) and a non-functional protein Bi-fp50x with the same molecular weight Bi-fp50 was generated.

The molecule weight of purified Bi-fp50 and the corresponding scFvs were first evaluated by sodium dodecyl sulfate-polyacrylamide gel electrophoresis (SDS-PAGE) with 15% gel (Fig. 1a). The Bi-fp50 (purity of 92% by ultraviolet analysis) had an approximate molecular weight of 50 kDa. At the same time, the two corresponding scFv was about 30 kDa, indicating that the fusion strategy could significantly decrease the molecular size if the length required to connect the two scFv were further considered. According to a previous analysis of protein diameter [31-32], the hydrodynamic diameter of the Bi-fp50 is about 5∼6 nm, which is very conducive to the penetration of the probe in the high-density PDAC tumors and timely renal clearing from the body [13]. Moreover, the obtained Bi-fp50 showed a typical protein 3D structure from the FTIR spectrum (Fig. 1b). It could be seen that the 1334 cm^-1^ and 1750 cm^-1^ represented the β-sheet and β-turn structure of the 3D protein, respectively. Furthermore, an enzyme-linked immunosorbent assay (ELISA) was carried out to measure the binding ability of the fusion protein Bi-fp50. As shown in Fig. 1c and d, Bi-fp50 showed a high target binding capacity comparable to the anti-VEGF scFv or anti-EGFR scFv, respectively. It was also noted that Bi-fp50 could recognize both targets simultaneously in a dose-dependent manner (Fig. 1e). The ELISA results reflected that the fusion protein Bi-fp50 had an excellent binding ability to VEGF and EGFR.

**Fig. 1.**
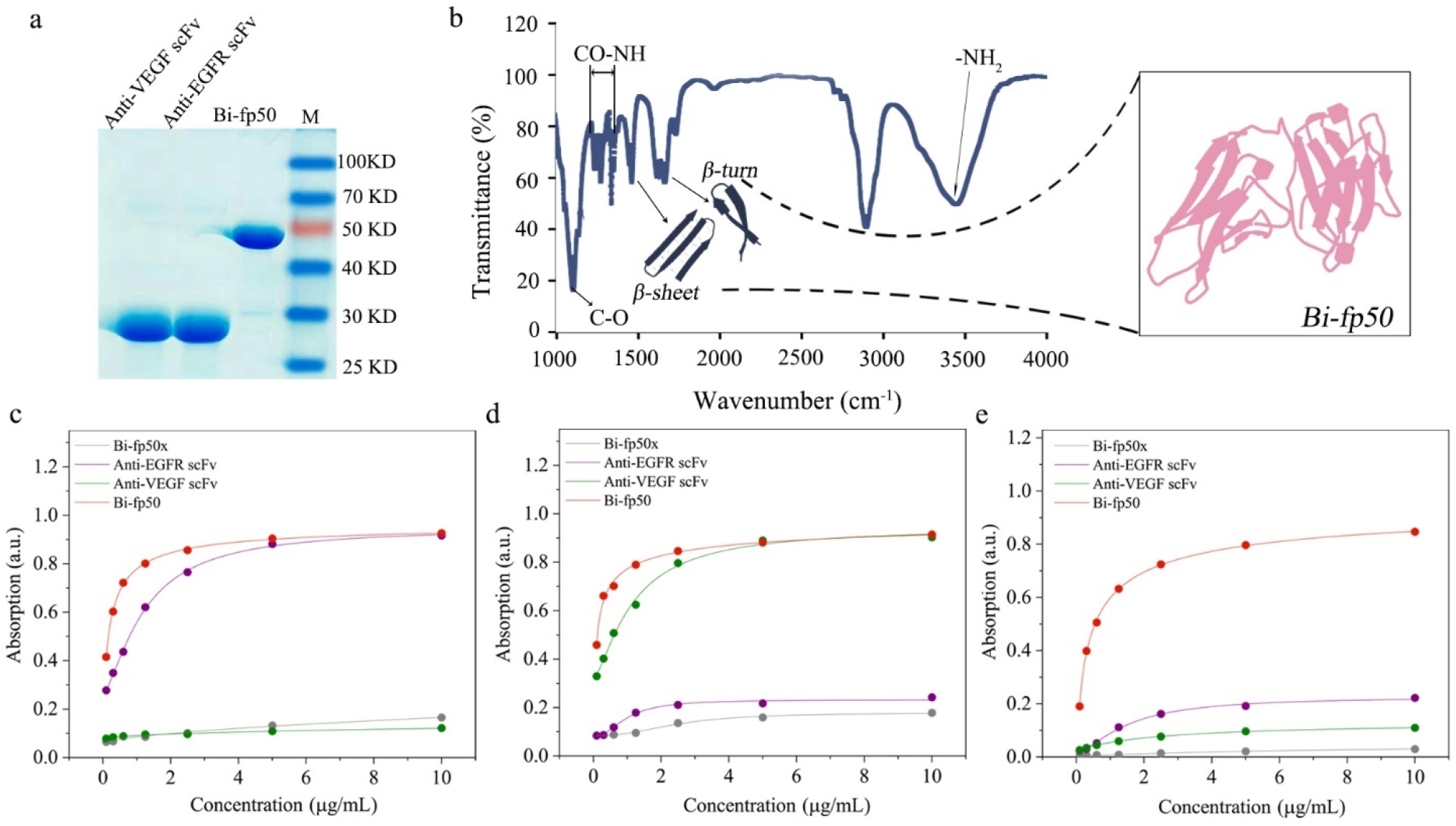
Characterization and binding affinity assessment of the Bi-fp50. **a**, SDS-PAGE analysis of the purified Bi-fp50 and two individual scFv. Bi-fp50 is about 50 kDa. **b**, Fourier-transform infrared spectroscopy (FTIR) spectrum of Bi-fp50, reflecting some typical structures and groups of the protein. **c-e**, The binding capacity of Bi-fp50 for EGFR (c), VEGF (d), EGFR, and VEGF simultaneously (e). Bi-fp50 shows a high affinity with human EGFR and VEGF.

### *In vitro* binding affinity of Bi-fp50

To quantitatively grasp the expression situation in PDAC, six typical PDAC cell lines were first chosen for the western blotting assay (Fig. 2a). And the anti-glyceraldehyde 3-phosphate dehydrogenase (GAPDH) antibodies were used as a loading control. All 6 cell lines showed relative solid abundance corresponding to moderate/high EGFR and VEGF overexpression. Further quantitative analysis showed that two cell lines with relatively higher EGFR and VEGF expression were Bxpc3 and Aspc1 (Fig. 2 b and c), used in the subsequent investigation. The *In vitro* binding affinity of Bi-fp50 in Bxpc3 cells was then evaluated after anti-EGFR scFv, anti-VEGF scFv, scFv2, Bi-fp50, and Bi-fp50x were labeled with AF488 (green fluorescence) dyes for visualizing.

**Fig. 2.**
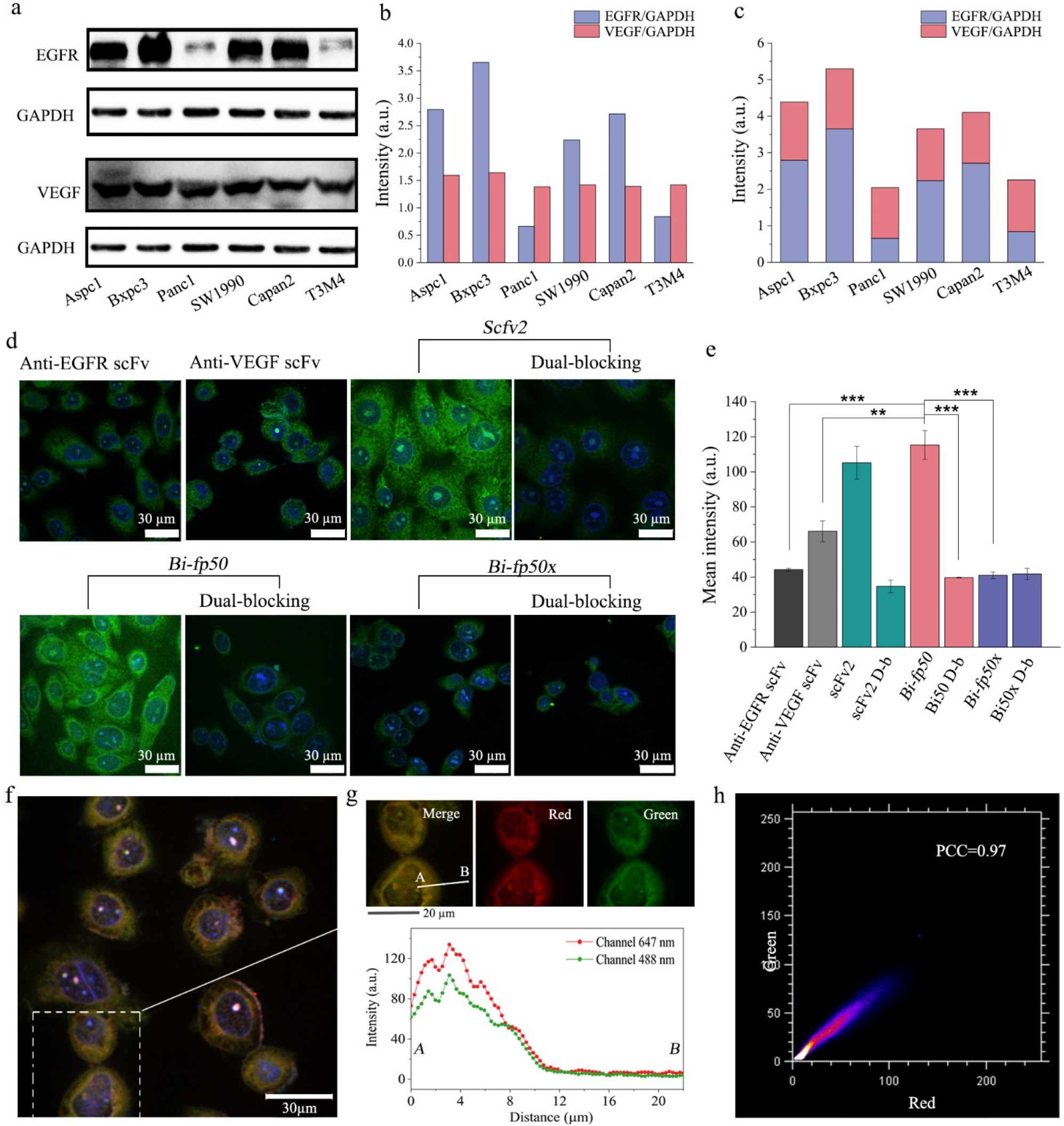
*In vitro* targeting assessment of Bi-fp50. **a**, Western blotting bands of EGFR and VEGF expression in different pancreatic cancer cell lines. glyceraldehyde-3-phosphate dehydrogenase (GAPDH) expression works as the standard inner line. **b-c**, Quantitative analysis of EGFR and VEGF expression individually (b) or accumulated (c) in six cell lines. **d**, Confocal images of Bxpc3 pancreatic cancer cells incubated with anti-EGFR scFv, anti-VEGF scFv, scFv2 (connected the two scFv), Bi-fp50, Bi-fp50x (the similar molecular weight of Bi-fp50, negative control). The Alexa Fluor 488 dye (AF488) was conjugated to image the probes and corresponding dual-blocking as the control groups. **e**, Quantitative analysis of the confocal images. Data are presented as mean ± standard deviation (SD); ** P<0.01, *** P<0.001. **f**, The confocal image of Bxpc3 cells was simultaneously incubated with anti-EGFR scFv-AF647 (red) and anti-VEGF scFv-AF488 (green). **g-h**, Location analysis of simultaneous targeting of EGFR (red) and VEGF (green) in a single cell (g) and corresponding colocalization coefficient (h), indicating the colocalization of VEGF and EGFR on the Bxpc3 cell surfaces.

As shown in Fig. 2d, two scFvs (anti-EGFR scFv, anti-VEGF scFv) had an apparent affinity for Bxpc3 cancer cells. ScFv2 group showed more vigorous fluorescence intensity with the Bxpc3 cells than each individual scFv, indicating an enhanced targeting effect. Similarly, the Bi-fp50 group had a significantly stronger green signal compared with single scFv groups, while the dual-blocking group had almost no fluorescence. Also, no green signal was observed in the Bi-fp50x and corresponding dual-blocking group. Quantitative fluorescence analysis (Fig. 2e) further reflected that the scFv2 and Bi-fp50 groups had significantly stronger fluorescence than the separately anti-VEGF scFv and anti-EGFR scFv groups. The fluorescence intensity of scFv2 and Bi-fp50 was approximately equal to the sum of that of two individuals. These results proved the enhanced, accumulated binding capability of Bi-fp50 and scFv for PDAC cells *in vitro*. To further explore the location of two scFvs (or Bi-fp50) bind cells, anti-EGFR scFv-AF647 (red) and anti-VEGF scFv-AF488 (green) were incubated with Bxpc3 cells simultaneously; two different fluorescence were found matched (Fig. 2f), suggesting the colocalization of two targets on the Bxpc3 cell surface. Additionally, the same pathway of two fluorescence channels from the selected area (Fig. 2g) and high Pearson’s r (PCC) of 0.97 (Fig. 2h) further demonstrated the excellent colocalization of VEGF and EGFR in Bxpc3 cancer cells. The obtained Bi-fp50 showed an excellent binding affinity for EGFR and VEGF with good colocalization performance in Bxpc3 cells.

### *In vitro* anticancer activity of Bi-fp50

Encouraged by the excellent binding ability for PDAC cells, the anticancer activity of Bi-fp50 for PDAC cells was further investigated *in vitro*. The Live/Dead staining assay intuitively and qualitatively showed that more Bxpc3 cells were dead in Bi-fp50 and scFv2 groups than in anti-EGFR scFv, anti-VEGF scFv, scFv2, and Bi-fp50x groups under the same concentration and incubation time (Fig. 3a). The cell viabilities of HPDE6-C7 normal and PDAC cells after incubated with anti-EFGR scFv, anti-VEGF scFv, and Bi-fp50 were quantitatively evaluated by MTT assay (Fig. 3b, c, and d). Using untreated cells as a normalized control, anti-EFGR scFv, anti-VEGF scFv, and Bi-fp50 treated HPDE6-C7 cells, human standard pancreatic duct epithelial cells with average level expression of EGFR and VEGF showed only a slight effect on growth even at high concentrations (Fig. 3b). Therefore, Bi-fp50 is relatively safe for normal cells at a certain concentration. By contrast, Bi-fp50 significantly inhibited the proliferation and growth of Bxpc3 and Aspc1 PDAC cells even under a relatively low concentration of 0.3 µM (Fig. 3c and d), while anti-EFGR scFv and anti-VEGF scFv alone had only a relatively slight inhibitory effect at the same concentration. The half-maximal inhibitory concentration (IC50) of Bi-fp50 to Bxpc3 and Aspc1 cells were about 0.32 µM and 0.38 µM, respectively, while the IC50 of two individual scFvs to PDAC cells was more than 1.0 µM (Fig. 3c and d). Therefore, we have the reason to believe that the fusion bispecific Bi-fp50 has a synergistically enhanced therapeutic effect relative to two individual scFv. The cell apoptosis was further studied by flow cytometry, as shown in Fig. 3e. It showed that Bi-fp50 treatment had significantly induced the early and late apoptotic of Bxpc3 cells. The early and late apoptosis together induced by Bi-fp50 was 75.7%, while 72.6% of scFv2, 67.7% of anti-EGFR scFv, and 55.7% of anti-VEGF scFv, respectively. Although anti-EGFR scFv and anti-VEGF scFv treatment could inhibit Bxpc3 cells, owing to the overexpression of EGFR and VEGF of Bxpc3 cells, they needed a longer time (48 hrs.), reflecting the low therapeutic effect. All these *in vitro* inhibition results showed the excellent therapeutic potential of Bi-fp50 for PDAC cells.

**Fig. 3.**
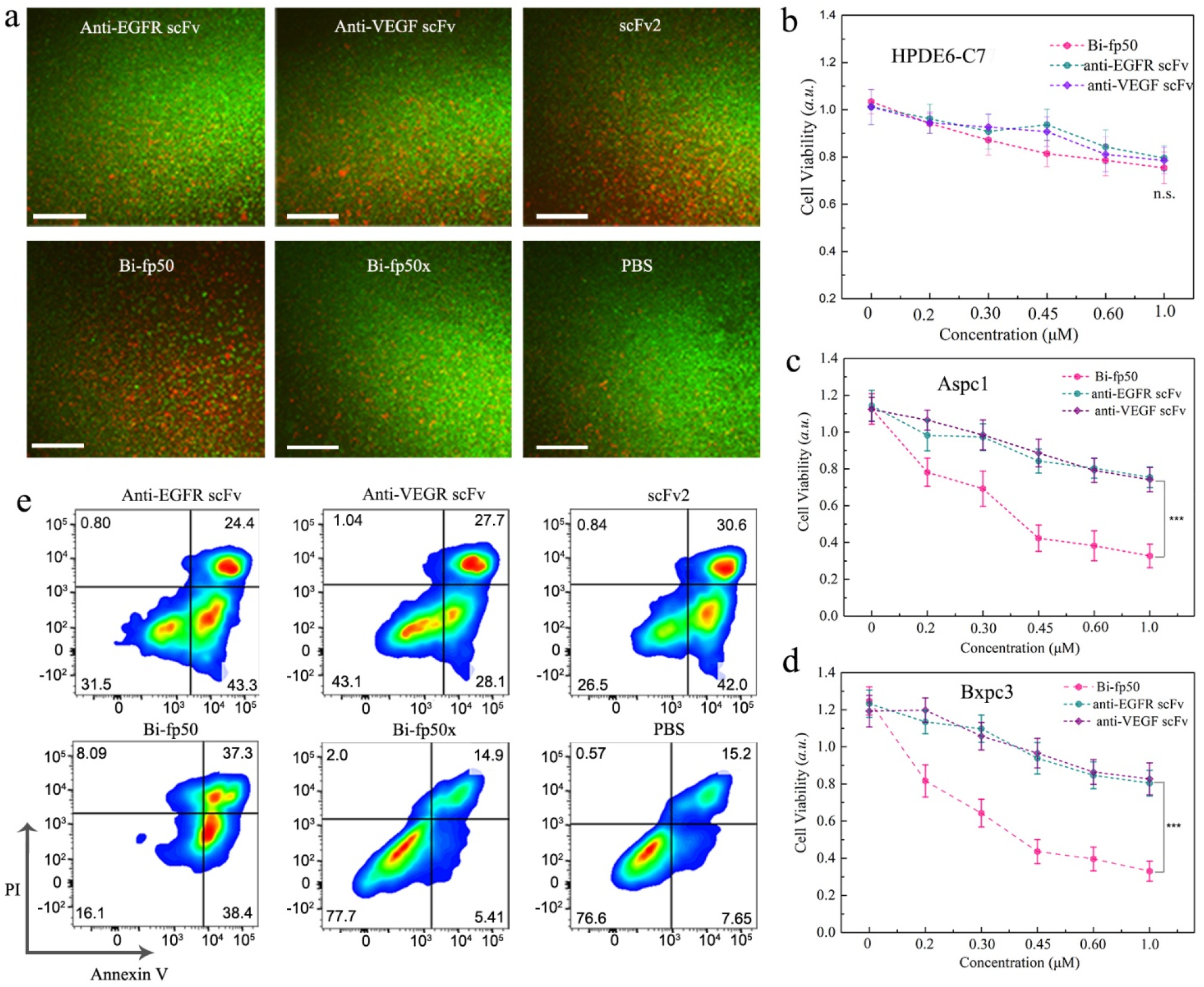
*In vitro* anticancer activity of Bi-fp50. **a**, Fluorescence images of Bxpc3 with different agents under the same concentration and time treatment (24 hrs.) stained with Calcein AM (live cells, green) and BOBO-3 Iodide (dead cells, red). Scale bars, 20 μm. **b-d**, Relative viability of Human Pancreatic Duct Epithelial Cell HPDE6-C7 (b), Pancreatic cancer cell Aspc1 (c) and Bxpc3 cells (d) after co-incubated with anti-EGFR scFv, anti-VEGF scFc, and Bi-fp50 for 24 hours. **e**, assessment of cellular death (apoptosis and necrosis) using Annexin V-FITC/PI staining after incubation with different probes of the same concentration for 48 h.

### Blood clearance, biodistribution, and *in vivo* tumor penetration of Bi-fp50

Before being applied *in vivo*, the pharmacokinetics and biodistribution of Bi-fp50 protein were first investigated. To visualize the probes, the near-infrared fluorescent molecule IRDye 800CW (Dye800) was conjugated with Bi-fp50 or scFv2 (Fig. S2). The Bi-fp50-Dye800 or scFv2-Dye800 (control group) were intravenously injected into orthotopic Bxpc3 tumor-bearing mice. The pharmacokinetics profile of the Bi-fp50-Dye800 was examined by fluorometry to determine the concentrations in blood at different time intervals post-injection (Fig. 4a). The half-life of blood clearance of Bi-fp50 was 4.33 ± 0.23 hours, and the area under the concentration-time curve (AUC) was 115 ID% h g^−1^ (Fig. 4a). This is like the pharmacokinetic profile of small-sized protein molecules (40∼50 KDa) [32]. The *in vivo* dynamic biodistribution of the Bi-fp50-Dye800 in the mice was then directly observed by fluorescence imaging (Fig. 4b). Considerable fluorescence of Bi-fp50-Dye800 in PDAC tumor could be observed from 2 h post-injection and gradually reached its maximum at 12 h, indicating the continuous accumulation of Bi-fp50-Dye800. *Ex vivo* fluorescence images of 12 h post-injection further conformed to the significant enrichment of Bi-fp50-Dye800 in the tumor. A quantitative analysis of the fluorescence signal to background ratio (SBR) of the tumor showed that the SBR of Bi-fp50 reached 3.65 post-injection of 4 hours (Fig. 4c) and continually increased to 3.73 in 8 hours and maintained a higher value to 12 hours, indicating the increased accumulation of probe in the tumor. In contrast, although the metabolic characteristics of scFv2-Dye800 were like those of Bi-fp50-Dye800, the enrichment in the tumor was intuitively lower than that of Bi-fp50-Dye800 at different times points (Fig. 4b and c). The AUC (area under the curve) of SBR was 87.99 h for Bi-fp50-Dye800, near two times of scFv2-Dye800 (45.26 h). Furthermore, a quantitative biodistribution analysis of major metabolic organs was conducted and further confirmed the higher tumor accumulation of Bi-fp50-Dye800 (Fig. 4d and e). Moreover, the Bi-fp50-Dye800 probes tended to enrich the kidney (4 h to 8 h) and then were metabolized mainly from the kidney, while the scFv2-Dye800 showed higher accumulation in the liver (Fig. 4d and e), owing to the effect of protein size. The faster clearance of Bi-fp50 from the body through the kidneys makes it more suitable for *in vivo* imaging and therapy, especially given the long-term safety profile [13, 32].

**Fig. 4.**
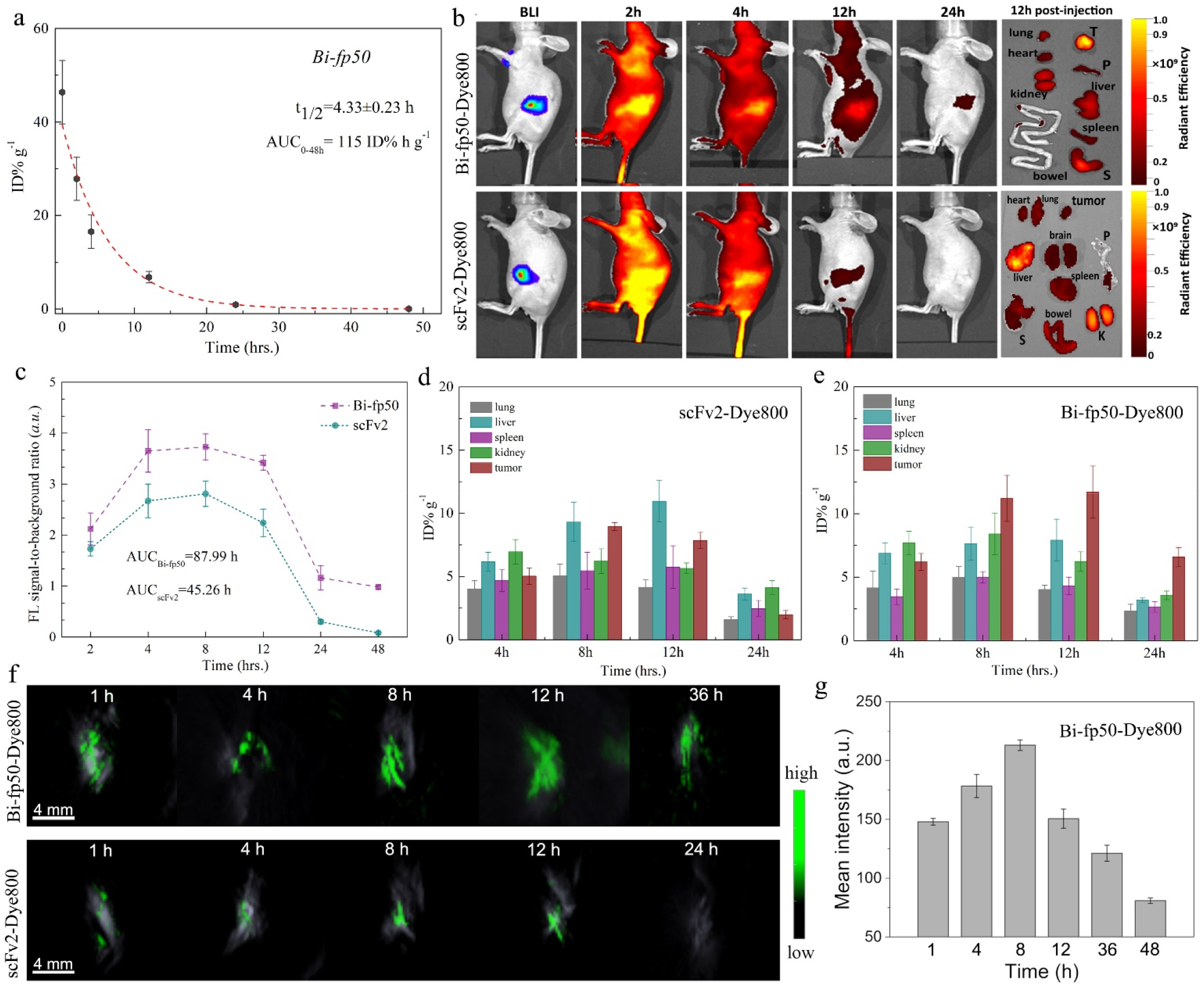
Pharmacokinetic and biodistribution evaluation of Bi-fp50. **a**, Blood circulation curve of the Bi-fp50-Dye800 determined by measuring the dye800 fluorescence intensity in the blood of the orthotopic Bxpc3 tumor-bearing mice at different time points post intravenous injection (n=5). **b**, *In vivo* fluorescence images of the Bi-fp50-Dye800 and scFv-Dye800-treated mice at different time points post intravenous injection. 12 hours after injection, tumors and major organs were harvested for *ex vivo* imaging. S, stomach; P, pancreas; T, tumor. **c**, Quantitative analysis of fluorescence intensity of Dye800 for Bi-fp50 and scFv2 in the tumor at different time points (n=5), the area of the small trapezoid under the curve is accumulated as AUC. **d-e**, Quantitative biodistribution analysis of the scFv-Dye800 (d) and Bi-fp50-Dye800 (e) in mice by measuring the Dye800 fluorescence intensity in the tumors and major organs at different times points post-injection. **f**, 3D photoacoustic images of orthotopic Bxpc3 tumors were acquired at different time points after injection of Bi-fp50-Dye800 and scFv-Dye800 probes. The bright signals are from the blood vessels in the tumor owing to hemoglobin. The green signal corresponds to the distribution of probes. **g**, Quantitative analysis of PAI intensity of tumor at different time points (n=3).

To further evaluate and confirm the dynamic penetration process of Bi-fp50 in the tumor, photoacoustic imaging (PAI) of the 3D model was carried out for orthotopic Bxpc3 tumor (Fig. S3). As a result, the Bi-fp50-Dye800 gradually penetrated the surrounding tumor parenchyma from local tumor vasculature (Fig. 4f). In contrast, the PAI signal of scFv2-Dye800 protein was much weaker than that of Bi-fp50-Dye800 at all the time intervals. And the signals of Bi-fp50 reached their peak in 8 hours (Fig. 4g), which was consistent with the quantitative results of fluorescence images. Additionally, PAI images of the 2D model (Fig. S4) showed a clear tumor profile by the Bi-fp50 probe in the tumor section, while only blurred boundaries could be imaged under the scFv2-Dye800 probe. Therefore, real-time *in vivo* imaging proved that the ultrasmall-sized fusion protein Bi-fp50 had a deep tumor tissue penetration compared with relatively large-sized scFv2.

### *In vivo* real-time tracking and locating Bi-fp50 in local pancreatic tumor

We further evaluated the dynamic and precise biodistribution of systemically administered Bi-fp50 in tumor microenvironments *in vivo*, which is believed to be vital for assessing medicines’ efficacy [33-34]. Fibered confocal fluorescence microscopy (CFL) contained two kinds of laser probes and was first used to visualize Bi-fp50 on a microscopic level in an orthotopic Bxpc3 pancreatic tumor model. The Bi-fp50 and scFv2 (control group) were labeled with AF647 (red) for real-time visualizing and imaging. Before injection of Bi-fp50 or scFv2, the normal pancreatic tissues, PDAC tumor, and junction of two different tissues could be distinguished and located very clearly after being stained by FITC (fluorescein isothiocyanate, green) dyes (Fig. 5a). Immediately after intravenous injection of Bi-fp50-AF647 or scFv2-AF647, more Bi-fp50 was enriched in tumor blood vessels than scFv2 (Fig. 5b). Furthermore, 8 hours post-injection, Bi-fp50 was widely distributed throughout the deep tissue of the tumor (Fig. 5c and Fig. S5), while only a small amount of fluorescence signal was observed in adjacent normal tissue and junction tissue. In contrast, only a tiny part of scFv2 was distributed in the deep tumor tissue, and a large proportion of scFv2 was concentrated in normal or junction areas (Fig. 5c and Fig. S5). The quantitative analysis of fluorescence intensity from CFL images confirmed such results and further revealed that the intensity of Bi-fp50 was about 2.5 times scFv2 in the deep tumor region and 1.7 times in the tumor of junction region (Fig. 5d). In addition, immunofluorescence (IF) images of the whole tumor section showed that the Bi-fp50 had penetrated the deep tumor tissue and distributed very uniformly across the entire tumor, while scFv2 was mainly spread in the surface layer of the rich vasculature (Fig. 5e).

**Fig. 5.**
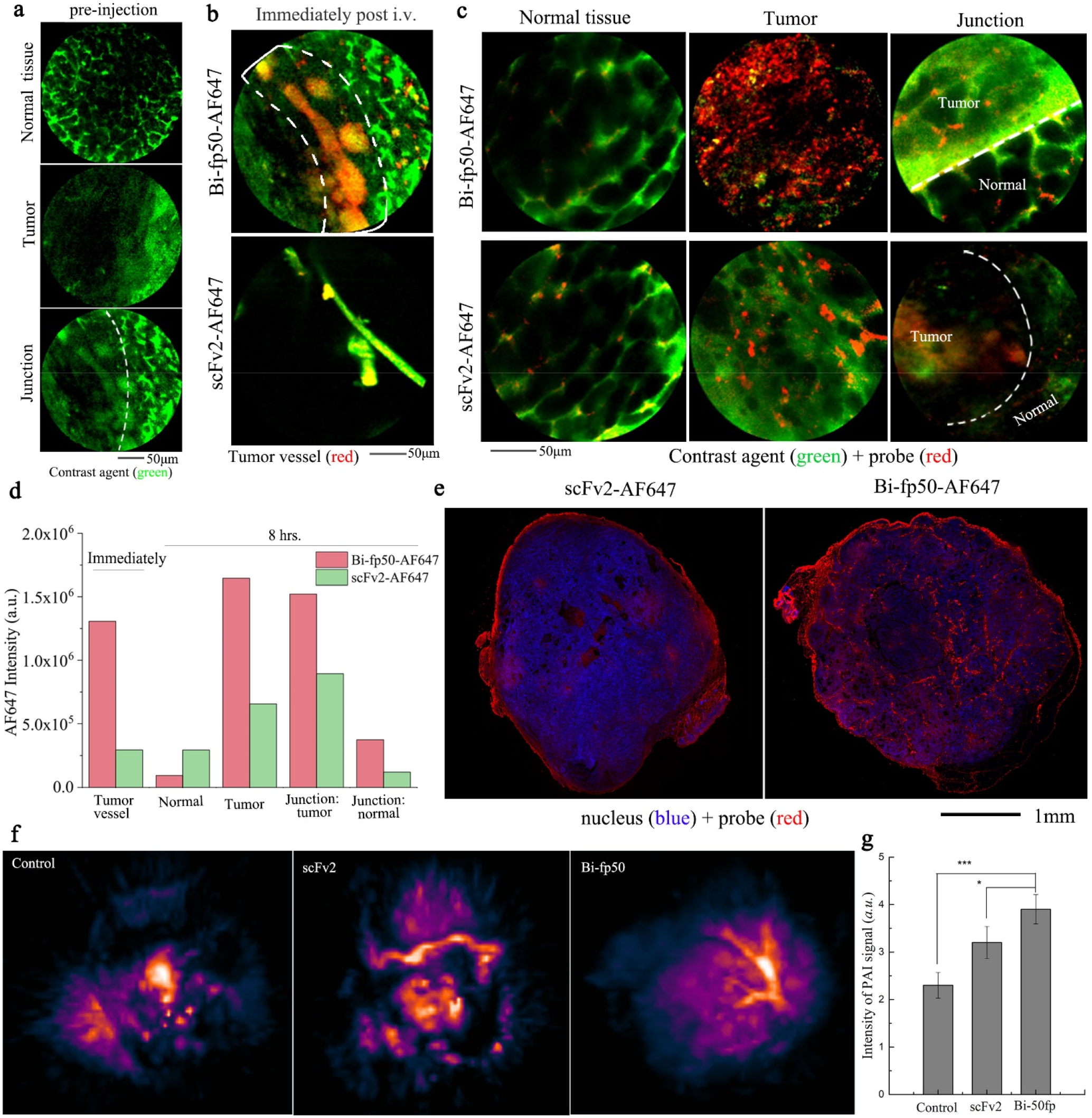
*In vivo* real-time, dynamic microscopical imaging of Bi-fp50 and scFv2 in local tumor of the orthotopic Bxpc3 tumor-bearing mice. **a**, Confocal fluorescence laser-microscopy (CFL) images of tumor tissue, surrounding normal tissue, and the junction between them pre-injection of probes. Single FITC dyes (green) were injected for 30 min before imaging to visualize the outline of tissues. **b**, CFL images of tumor tissue immediately post i.v. Injection of probes (red). **c**, CFL images of tumor, normal and junction tissues 8 hours post i.v. Injection of probes (red). **d**, Quantitative analysis of fluorescence intensity of Bi-fp50 or scFv2 in different positions of PDAC tumor at different time points. e, Immunofluorescence (IF) images of the whole tumor tissues resected 8 h after intravenous injection of scFv2 or Bi-fp50. The probes were labeled with AF647 (red), and the nuclei were stained with 4’,6-diamidino-2-phenylindole (DAPI). f, 3D photoacoustic images of orthotopic Bxpc3 tumors were acquired 12 hours after different injections: PBS (control), scFv2, and Bi-fp50. The bright signals are from the blood vessels in the tumor owing to hemoglobin. Brightness is positively correlated with hemoglobin content and identifies the distribution of tumor micro-vessels. g, Quantitative analysis of PAI signals (n=3), * P<0.05, *** P<0.001.

In addition to targeting PDAC cells, Bi-fp50 can also act on the tumor’s vasculature by targeting the VEGF pathway to improve tumor vessel perfusion [26-27]. The 3D PAI images of the orthotopic Bxpc3 pancreatic tumor showed that compared with the control group (non-treated) and the group injected with scFv2, the tumor vasculature of the Bi-fp50-treated group became significantly more abundant, and the blood flow also increased 12 hours post-injection (Fig. 5f). The quantitative analysis of PAI signals further confirmed that the improvement of tumor vessel perfusion of Bi-fp50 was more significant than the bispecific scFv2 (Fig. 5g). The ultra-small size of Bi-fp50 allows it to penetrate deeper tissues, thereby altering the broader vascular system. The increase of vascular permeability can, in turn, promote the penetration of Bi-fp50. Thus, the interaction of ultra-small size and anti-VEGF capability allows for better penetration and distribution of Bi-fp50. The significant enrichment and uniform distribution of Bi-fp50 in tumor tissues, especially the penetration into deep tissues, lays the foundation for the exertion of bispecific therapeutic effects.

### *In vivo* antitumor efficacy of Bi-fp50

Encouraged by promising results of *in vitro* therapy and *in vivo* imaging, we further explored the antitumor efficacy of Bi-fp50 *in vivo*. 10 Balb/c nude mice with Bxpc3 xenograft pancreatic tumors were randomly divided into two groups (n=5). After intravenously injected with Bi-fp50 (10 mg Kg^-1^) or scFv2 (10 mg Kg^-1^), tumor size and the bodyweight of mice were continually recorded every 2 days. The substrate to activate Luc-protein of Bxpc3 cells was administered through intraperitoneal injection to visualize the tumor size (Fig. 6a). After 3 weeks of treatment, the bioluminescence imaging (BLI) and the representative photographs of tumors showed that the tumor size in the Bi-fp50 group was smaller than that in the scFv2 group (Fig. 6a and c). In addition, the Bi-fp50 group showed a higher antitumor efficacy with a tumor inhibition ratio of 35.14% compared to the scFv2 group as a control (Fig. 6b). The weight of the two groups of mice remained stable, and there was no significant difference (Fig. 6d). The anticancer efficiency was further evaluated by a terminal deoxynucleotidyl transferase-mediated deoxyuridine triphosphate nick end-labeling (TUNEL) assay, which is widely utilized to study the intra-tumoral apoptosis after therapy treatment [references]. As shown in Fig. 6e, few TUNEL-positive cells (green) were observed from the scFv2-treated tumor section. In contrast, many TUNEL-positive apoptotic cells (shown in green) could be kept in the Bi-fp50-treated group, distributed throughout the entire tumor section. These results indicate that the Bi-fp50 can induce PDAC cell death by activating apoptosis in the tumor. 3 weeks after different treatments, mice were sacrificed, and major organs (heart, liver, spleen, lung, kidney, and pancreas) were harvested for histological evaluation. The H&E staining showed no significant abnormalities, and differences in major organs were observed for Bi-fp50x (control) and Bi-fp50 groups (Fig. 6f). These results indicated the Bi-fp50 protein could significantly inhibit tumor growth by intravenous injection *in vivo* without other noticeable side effects. After investigating the antitumor effect of Bi-fp50, the molecular mechanism underlying this effect was tentatively explored. We detected the activation of downstream molecules of EGFR and VEGF. In Bxpc3 cells, the constitutive phosphorylations of STAT3 and AKT signal pathways (P-STAT3, P-AKT) were more strongly inhibited by Bi-fp50 than by other agents (scFv2, anti-EGFR scFv, anti-VEGF scFv, and Bi-fp50x) as shown in Fig. 6g and Fig. 6h. It has been reported that the targeting of EGFR and VEGF of PDAC cells will affect cell survival mainly through STAT3 and AKT pathways [13, 20]. Furthermore, inhibition of the expression of EGFR and downstream signaling pathways can induce apoptosis of pancreatic cancer cells, and downstream signaling pathways related to VEGF can affect cancer angiogenesis [35]. Therefore, simultaneous binding of Bi-fp50 to EGFR and VEGF can cross-talk and regulate AKT- and STAT3-related pathways together, thereby affecting the cell growth, survival, angiogenesis, and migration of PDAC cells (Fig. 6i).

**Fig. 6.**
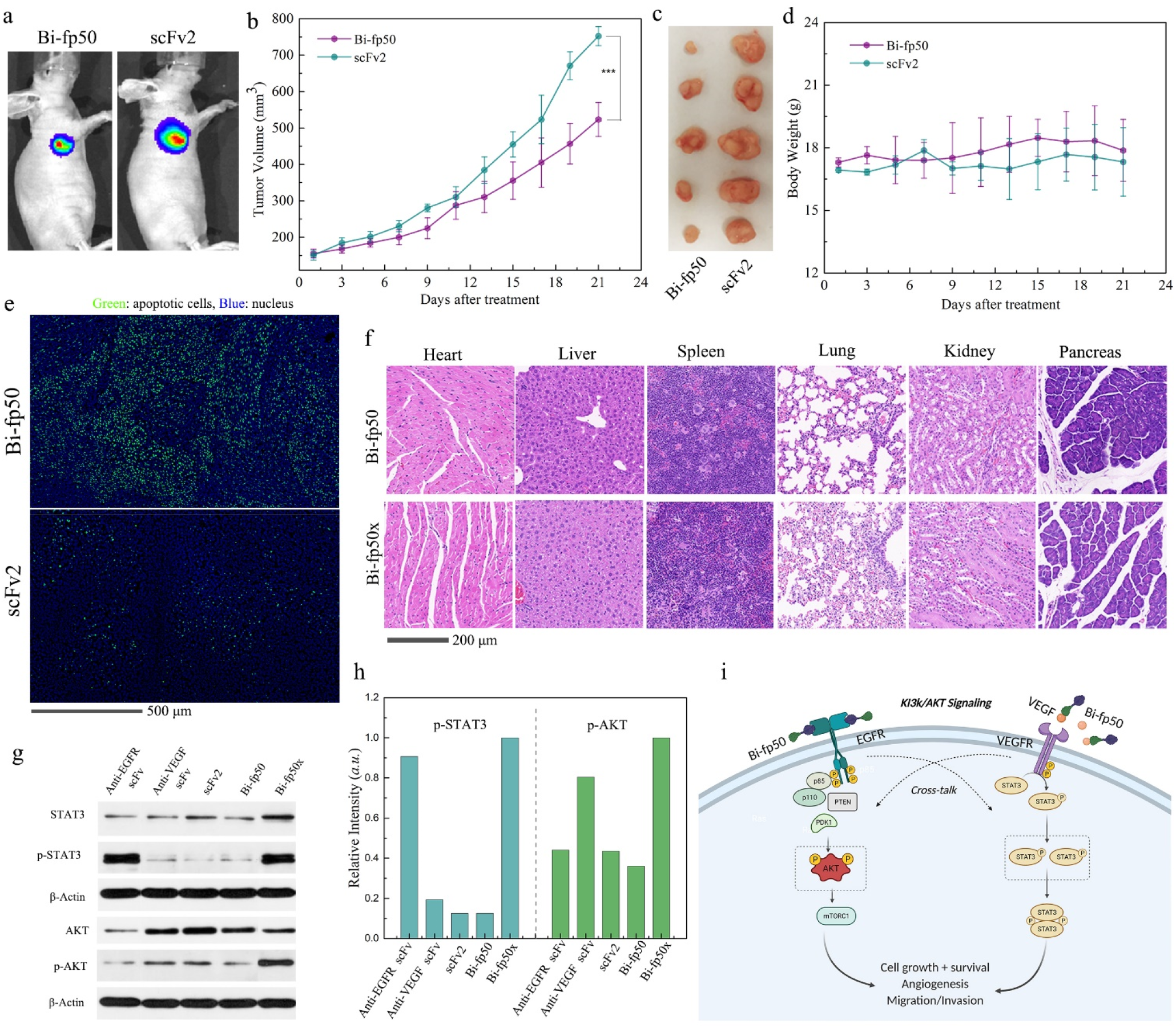
*In vivo* evaluation of antitumor efficacy of Bi-fp50. **a**, Bioluminescence imaging (BLI) of xenograft Bxpc3 tumor-bearing mice 3 weeks after different treatment. **b**, Growth curves of Bxpc3 tumors in different groups of nude mice treated with Bi-fp50 or scFv2 with a dose of 10 mg kg^−1^. Data are mean ± s.d. of biological replicates (n = 5). *** *p* < 0.001. **c**, tumors were dissected on day 22 for the Bi-fp50 and scFv2 groups. **d**, body weights of mice in different groups 3 weeks after treatments. Data are mean ± s.d. of biological replicates (n = 5). **e**, TUNEL (terminal deoxynucleotidyl transferase-mediated deoxyuridine triphosphate nick-end labeling)-stained tumor sections five days after different treatments. Apoptotic cells are shown in green. **f**, H&E staining of major organs 3 weeks after Bi-fp50 and Bi-fp50x (negative control) treatment. No differences were found between the two groups. **g**, Western blotting bands of STAT3 and AKT signal pathway. β-Actin was the standard inner line. **h**, Relative intensity of p-STAT3 and p-AKT proteins. **i**, Schematic illustration of main inhibition mechanism of Bi-fp50 in EGFR and VEGF signal for PDAC.

## Conclusion

We constructed the bispecific fusion protein Bi-fp50 that could target both EGFR and VEGF of PDAC cells simultaneously. Due to the cross-talk and inter-regulation of the EGFR-VEGF pathway, the Bi-fp50 achieved an enhanced, synergistic therapeutic effect in Bxpc3 and Aspc1 PDAC cells. Furthermore, under the combined impact of ultrasmall small size and function of tumor vasculature normalization, the Bi-fp50 largely penetrated dense barriers and enriched the whole Bxpc3 pancreatic tumor to achieve a significant tumor inhabitation effect *in vivo*. Therefore, the synthetic Bi-fp50 bispecific protein showed excellent potential as an efficient pathway-targeted specific therapy for PDAC used alone or in combination with other therapeutic modalities.

Multi-specific biological molecules, especially bispecific antibodies and their derivatives scFvs, have rapidly grown clinic attention due to the unique advantages of providing off-the-shelf immunotherapeutics for treating cancer [36-38]. However, as with any immunotherapy, safety and off-target toxicity issues are a potential concern. Bispecific antibodies or derivatives that bridge T cell-cancer cells with two terminals of high-specific selectivity simultaneously can significantly decrease the risk of off-target side effects. Additionally, when this strategy is applied to solid tumors, especially very dense PDACs, the intratumoral enrichment and penetration, and timely clearance from the body are critical for multi-specific biological molecules. Therefore, the Bi-fp50, with its ultra-small size and multispecificity, also has excellent potential in PDAC dual-targeted immunotherapy.

## Authorship contribution statement

Q.W., H.Y., J.T., and X.M.Z. discusses and convinced this project. Q.W., J.Y.W., and H.Y. designed the experiments. Q.W. and Z.L. fabricated the Bi-fp50 protein. Q.W. and J.Y.W. performed the characterization, *in vitro* imaging, and therapy assessment. Q.W. developed the PDAC animal model. Q.W. and K.W. performed *in vivo* imaging experiments. Q.W., J.Y.W., and H.Y. performed *in vivo* therapy experiments. F.Y.K assisted in data analysis. Q. W., J.Y.W., and H.Y. prepared figures and wrote the manuscript, with revision by J.T. and X.M.Z.

## Acknowledgments

We thank the technical support from the Multimodal Biomedical Imaging lab, Institute of Automation, the Chinese Academy of Sciences, and the Cold Spring Biotech Co., Ltd. We thank Beijing Gegen Biotechnology Co., Ltd. for help in preparing and purifying the scFv fragments and fusion protein. This work was supported by the National Key Research and Development Program of China (grant nos. 2017YFA0205200) and the National Natural Science Foundation of China (grant nos. 81,901,813, 81,671,757, 92,159,305), and the CAMS Innovation Fund for Medical Sciences (grant no.2021-I2M-C&T-B-067).

## Supplementary data

Supplementary data to this article can be found online at https://

## Data availability statement

The data are available from the corresponding author on reasonable request.

## Conflict of interest

The authors declare no conflicts of interest.

## Notes

### Competing Interest Statement

The authors have declared no competing interest.

